# Efficient Genomic Control for Mixed Model Associations in Large-scale Population

**DOI:** 10.1101/2021.03.10.434745

**Authors:** Zhiyu Hao, Jin Gao, Yuxin Song, Runqing Yang, Di Liu

## Abstract

Among linear mixed model-based association methods, GRAMMAR has the lowest computing complexity for association tests, but it produces a high false-negative rate due to the deflation of test statistics for complex population structure. Here, we present an optimized GRAMMAR method by efficient genomic control, Optim-GRAMMAR, that estimates the phenotype residuals by regulating downward genomic heritability in the genomic best linear unbiased prediction. Even though using the fewer sampling markers to evaluate genomic relationship matrices and genomic controls, Optim-GRAMMAR retains a similar statistical power to the exact mixed model association analysis, which infers an extremely efficient approach to handle large-scale data. Moreover, joint association analysis significantly improved statistical power over existing methods.

## Introduction

The linear mixed model (LMM) ^1,2^ is an efficient and practical approach for genome-wide association analysis (GWAS) because of its ability to comprehensively control confounders, such as population stratification, family structure, and cryptic relatedness ^3^. The LMM-based association analysis estimates corresponding variance components to each single nucleotide polymorphism (SNP), followed by statistical inferences on the association of the SNP with a phenotype using the generalized least square method^4^, given a phenotypic covariance matrix. Imaginably, such a genome-wide mixed model association is compute-intensive when handling high-throughput SNPs.

Simplifying genome-wide mixed model association launched from reducing estimation for variance components with entire genomic markers ^5–8^, clustering subjects or sampling markers ^6,8–10^. For this reason, the spectral decomposition ^5^, the Monte Carlo REML ^11,12^ and the H-E regression ^13,14^ methods sequentially were considered as the alternative of maximum likelihood and restricted maximum likelihood (REML) ^15^. As a potential option, the Method R ^16,17^ is able to estimate the variance components in a computationally efficient manner by repeatedly solving the genomic best linear unbiased prediction (GBLUP) equations ^18^. Subsequently, several simplified algorithms were proposed to replace different polygenic effects or variances related to high-throughput markers with the same genomic breeding values (GBVs) or genomic variance ^9,19–21^. In genomic selection ^22–24^, many methods except for GBLUP can estimate GBVs, avoiding pre-estimation of variance components. In animal breeding ^25^, people always robustly evaluate variance components by drawing the balanced sample from a large population, this suggests that the genomic relationship matrix (GRM) ^26^ can also be calculated with sampling subjects for estimating variance components in genome-wide mixed model association. Transforming human GRM to the sparse, recently, a fastGWA ^27^ was developed upon GRAMMAR-Gamma to extreme efficiently achieve large-scale mixed model association analyses.

These simplified algorithms efficiently save computational costs in terms of building the GRM, estimating the variance components or computing the association statistics for each SNP. However, they do output a certain degree of statistical error due to under/over-estimation of the polygenic effects and heritability. Among the simplified algorithms, GRAMMAR ^19^ performs the lowest computing time of *O(nm)*, where *n* is population size and *m* is the number of markers, for the association tests. Based on the GRAMMAR algorithm, we regulated downward genomic heritability in GBLUP equations to precisely estimate the polygenic effects, correcting the genome-wide deflation of association statistics by optimizing the genomic control (GC). The optimized GRAMMAR algorithm, named Optim-GRAMMAR only requires to repeatedly solve the GBLUP equations, as BOLT-LMM ^8^ did in estimating variance components. Further, we performed joint analyses of multiple QTN candidates obtained with multiple testing, to significantly improve the statistical power to detect QTNs.

## Results

### Statistical properties of Optim-GRAMMAR

We recorded association results obtained with Optim-GRAMMAR, a test at once for the simulation (1), and compared them with FaST-LMM, GRAMMAR, GRAMMAR-Gamma, and BOLT-LMM. The Q-Q and ROC profiles are shown in Figure 1 and Figure 2 for the 200 simulated QTNs (Figure 1S and Figure 2S for simulation scenarios). Under the optimized GC that infinitely close to 1.0, Optim-GRAMMAR yielded almost the same statistical power to detect QTNs as the exact FaST-LMM. This was not affected by the number of QTNs or the genomic heritability simulated. As usual, statistical powers increased with the number of QTNs and the genomic heritability increased, no matter for how population complex. For the maize population, a complex population structure caused slightly lower statistical power when using Optim-GRAMMAR than with the FaST-LMM, though the difference was negligible. GRAMMAR had the lowest GC and statistical power across all the methods. Moreover, population structure was more complex, false negative rate was higher for GRAMMAR. Although GRAMMAR-Gamma and BOLT-LMM corrected the underestimated test statistics by GRAMMAR using the defined calibration factors ^8,21^, GRAMMAR-Gamma still slightly deflated the test statistics when dealing with complex maize population, generating a GC of less than 1.0. In contrast, BOLT-LMM, strongly increases the false positive rates in similar populations due to inflation of test statistics.

**Figure 1:**
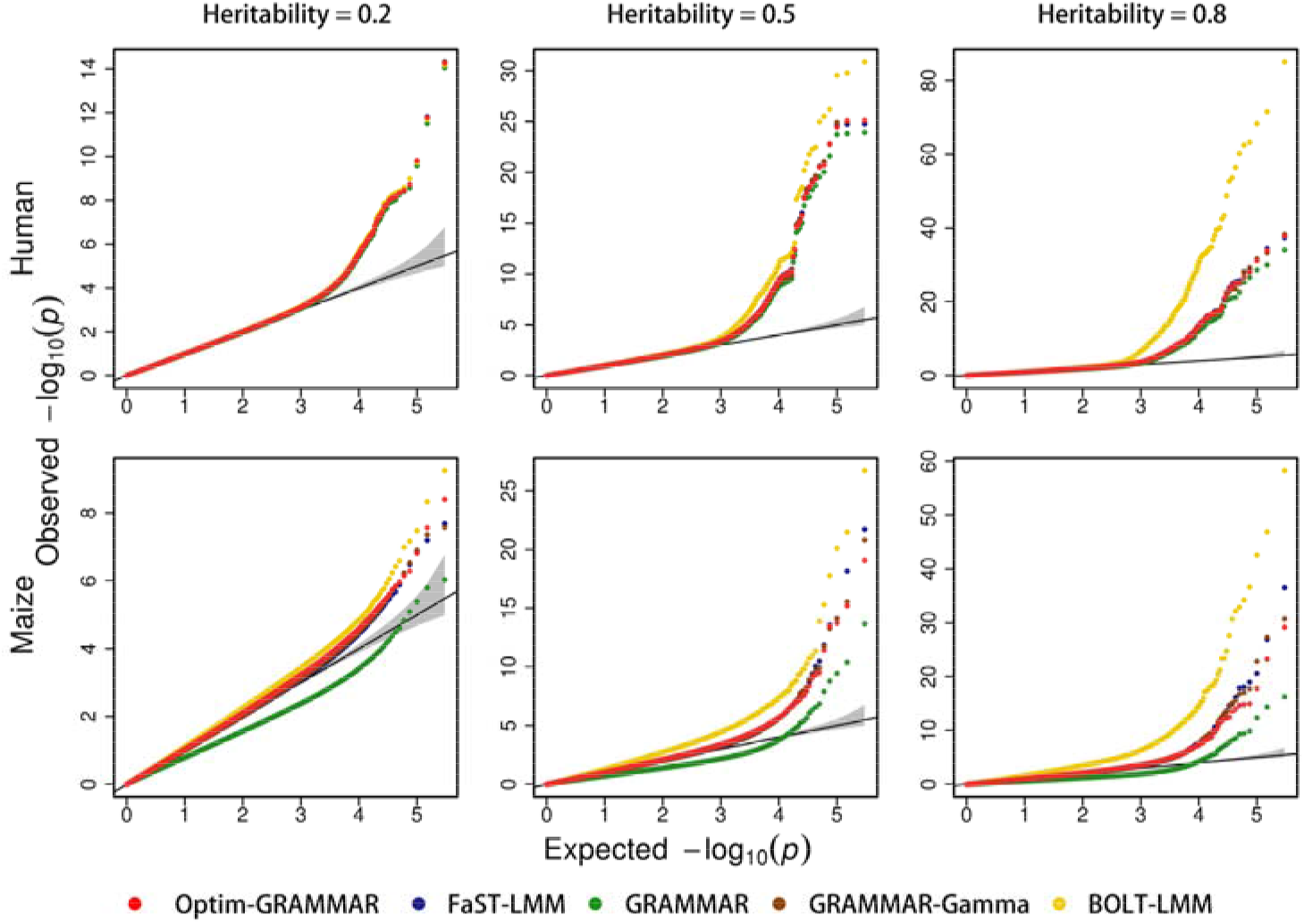
Comparison in the Q-Q plots between the Optim-GRAMMAR and the four competing methods. The simulated phenotypes are controlled by 200 QTNs with the low, moderate and high heritabilities in human and maize. The Q-Q plots for all simulated phenotypes are reported in Supplementary Figure 1S.

**Figure 2:**
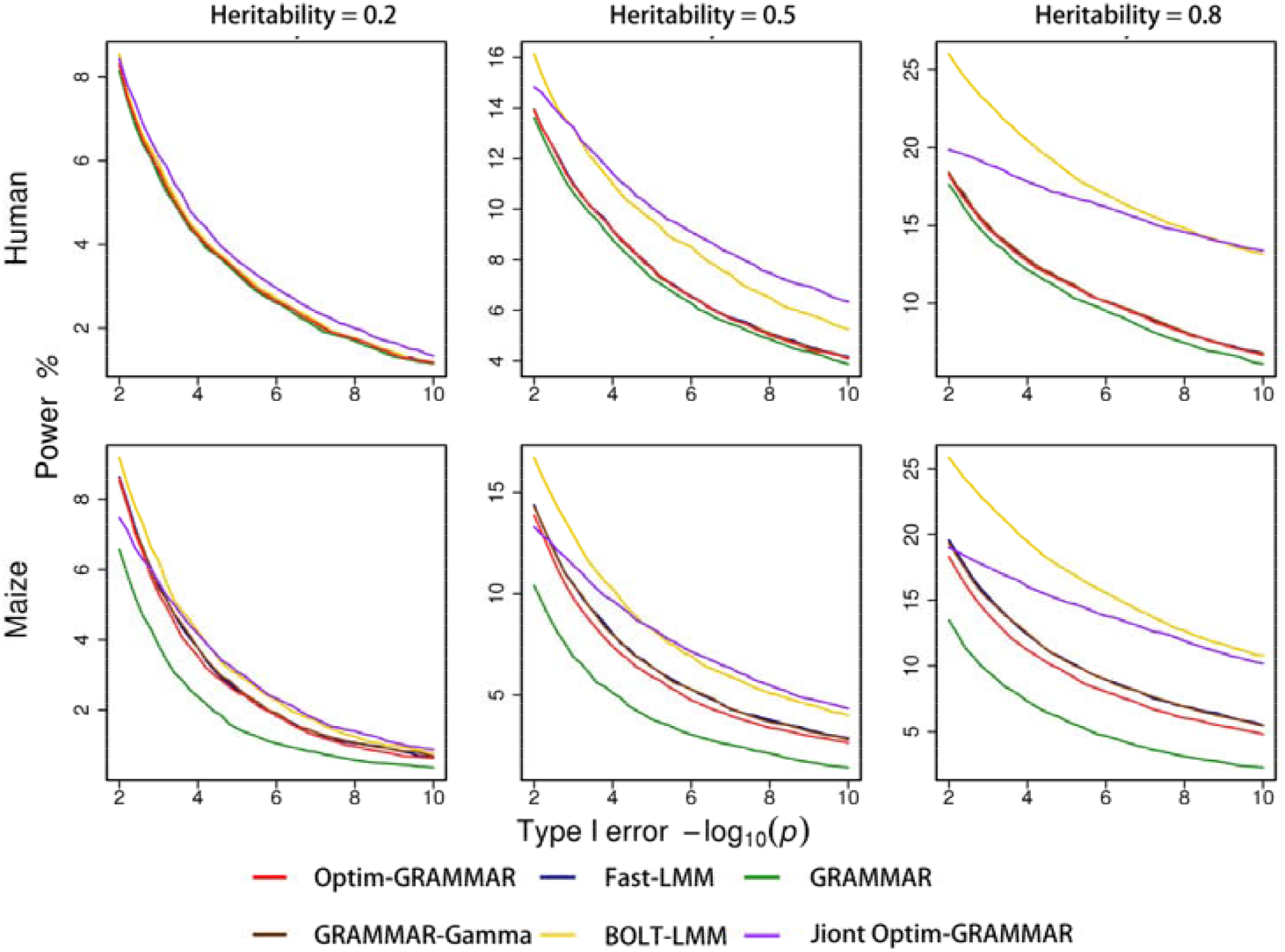
Comparison in the ROC profiles between the Optim-GRAMMAR with the four competing methods. The ROC profiles are plotted using the statistical powers to detect QTNs relative to the given series of Type I errors. The simulated phenotypes are controlled by 200 QTNs with the low, moderate and high heritabilities in human and maize. The ROC profiles for all simulated phenotypes are reported in Supplementary Figure 2S.

After optimizing the GC, Optim-GRAMMAR jointly analyzed multiple QTN candidates chosen from an association test at a time, at a significance level of 0.05. For convenience to compare, we depicted together statistical powers obtained with a test at once and joint association analyses. Using backward regression analyses, we demonstrated that Optim-GRAMMAR had increased statistical powers. In comparison, BOLT-LMM also had the higher statistical powers, but could not control the false-positive rates, specifically for complex population structure.

### Calculation of GRMs and GCs with the sampling markers

With the GBLUP, estimation of both genomic heritability and GBVs depends on the marker density used to calculate the GRM in the structured population ^26,28^. For simplification to compute, the FaST-LMM and BOLT-LMM tried to sample or screen a small proportion of the whole genomic SNPs to estimate the genomic heritability or GBVs as precisely as possible. In contrast, Optim-GRAMMAR would require the fewer sampling markers to assist the underestimation of GBVs, which are demonstrated by simulation (2).

Figure 3 showed the changes in GCs with numbers of sampling markers obtained with Optim-GRAMMAR and four competing methods. No competing method controlled for the false-positive or false-negative errors with less than 50,000 SNPs. Both FaST-LMM and GRAMMAR-Gamma gradually controlled the false-positive errors as the sampling markers increased. GRAMMAR seemed to overcome the false-negative errors by using less than 3000 sampling markers to underestimate the GBVs, whereas BOLT-LMM still yielded very high false-positive errors, despite using the sampling markers to estimate the GRM. As shown in Figure 3S and Figure 4S, Optim-GRAMMAR still retained a high power to detect QTNs through the optimization of the GCs, even using less than 3000 sampling markers. At high heritability simulated, little difference in statistical power happened across the clusters of sampling SNPs, this suggested that for Optim-GRAMMAR, the less sampling SNPs contributed to GC for the traits of lower-moderate heritability (See Supplementary Table 1S).

**Figure 3:**
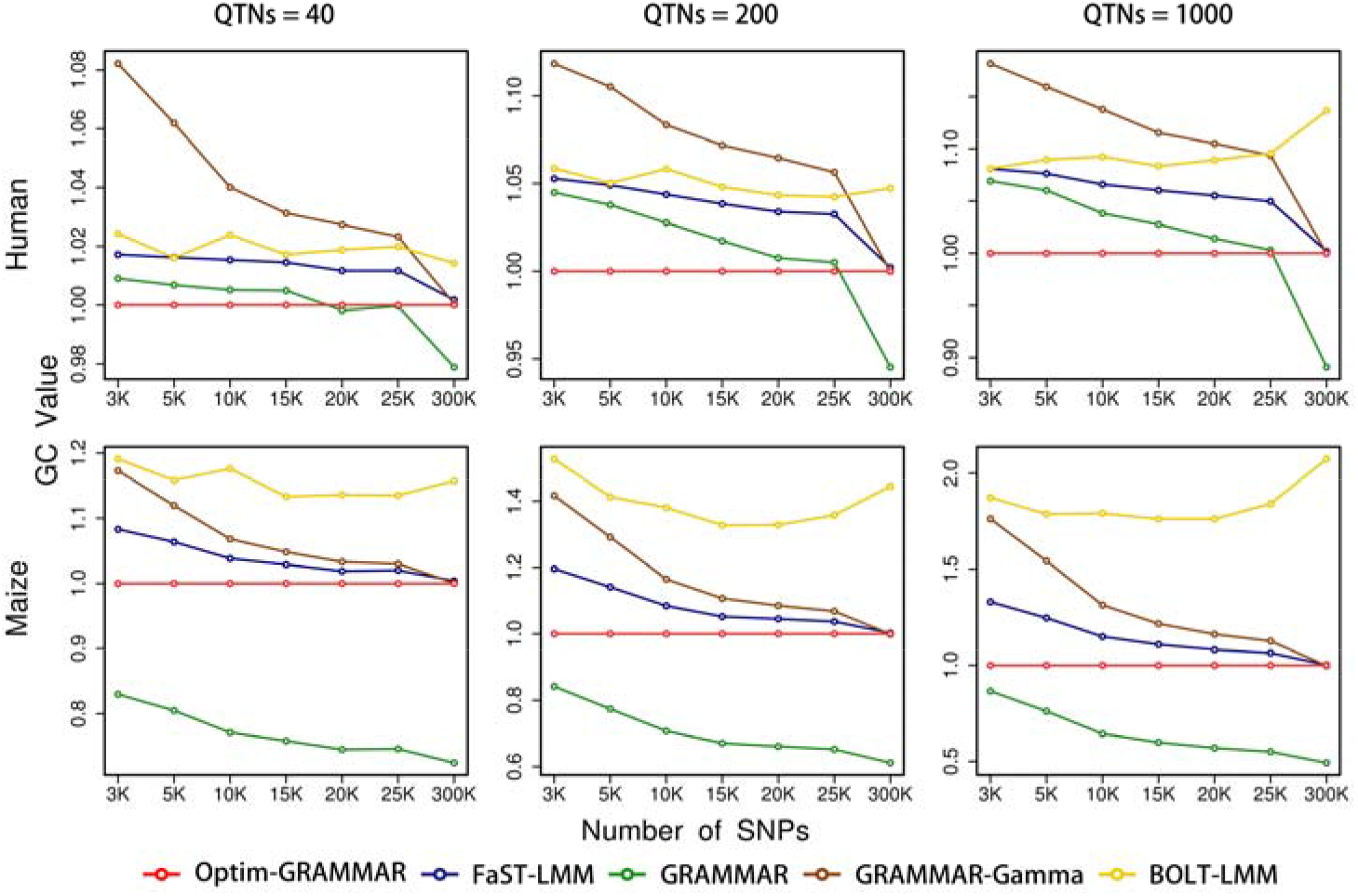
Changes in GCs with the number of sampling SNPs for calculating GRMs with the Optim-GRAMMAR and the four competing methods. GC is calculated by averaging genome-wide test statistics. The simulated phenotypes are controlled by 40, 200 and 1,000 QTNs with the moderate heritability in human and maize.

**Figure 4:**
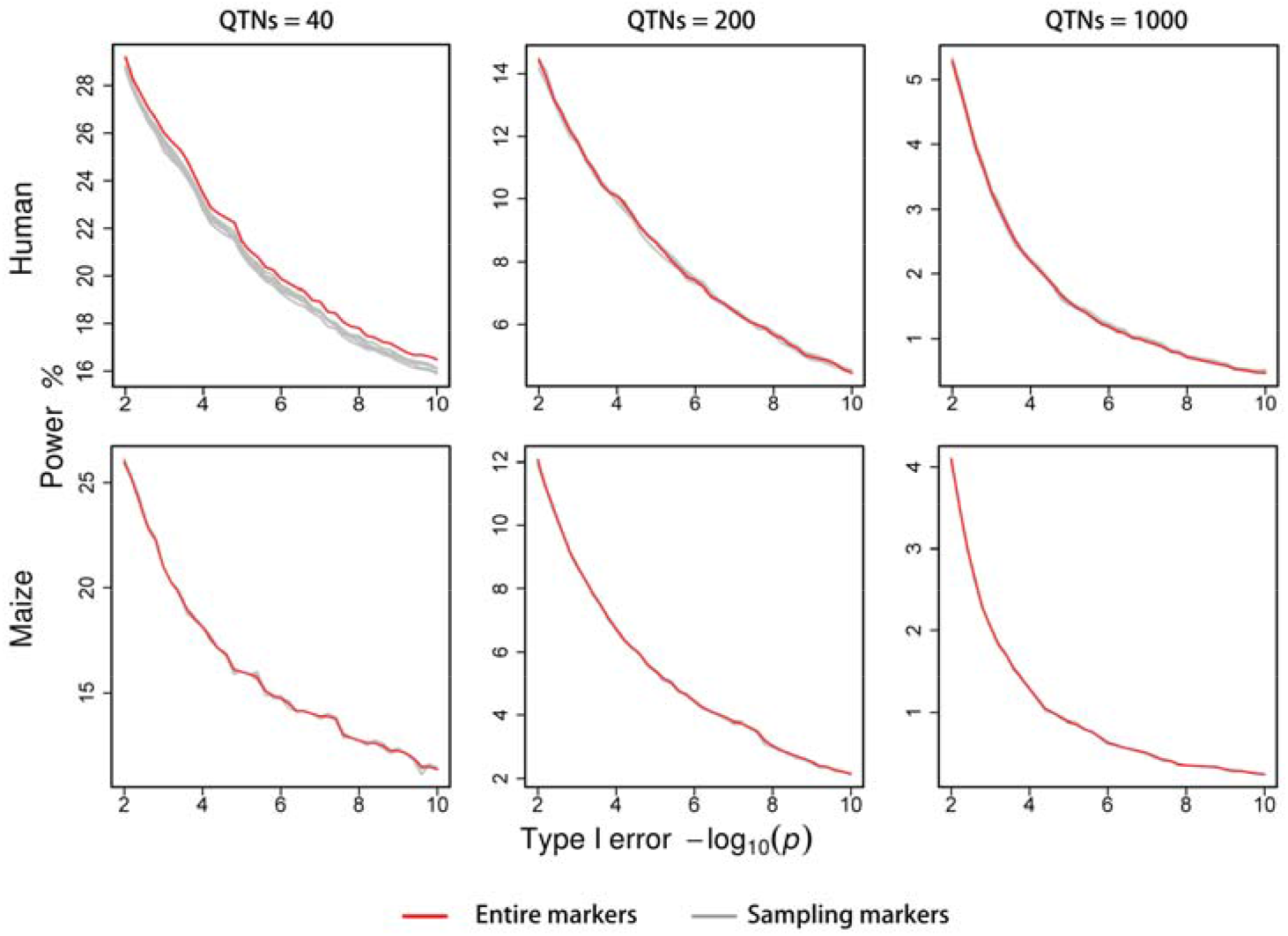
Changes in statistical powers with the number of sampling SNPs for calculating GCs with the Optim-GRAMMAR. Statistical powers are dynamically evaluated with the ROC profiles. The simulated phenotypes are controlled by 40, 200 and 1,000 QTNs with the moderate heritability in human and maize.

To further improve computing efficiency to optimize GCs, in updating genomic heritability we evaluated GCs by using different numbers of sampling markers under calculating GRMs using 5,000 sampling SNPs. The effects of the sampling markers on association results are summarized in Table 1S for GCs and GBVs, Figure 4 for ROC profiles and Figure 5S for Q-Q plots. In optimizing GRAMMAR, as expected, calculating GCs using more than 5,000 sampling SNPs could accurately estimate polygenic effects, ensuring high statistical powers in almost the same GCs as those using entire markers. This would extremely simplify mixed model association analysis for large-scale data.

### Real data analyses

Public datasets for the HS and CFW mice and maize were downloaded and analyzed with Optim-GRAMMAR to generate both QTN mapping and GC. We filtered out 109 from 973 phenotypes in the HS mouse and 36 from 123 phenotypes in the CFW mouse using a normality test. We only recorded the computing times for the maize dataset, because both of the mouse populations either had very small population sizes or small marker sample sizes that would not adequately show the difference in computing complexity.

The three datasets were analyzed using all of the competing methods previously used in our simulation experiments. The Q-Q and Manhattan profiles for the traits of detectable QTNs are shown in Supplementary Figure 4S (HS mouse), Figure 5S (CFW mouse), and Figure 6S (maize). As shown in the plots, GRAMMAR, GRAMMAR-Gamma, and BOLT-LMM behaved almost identically in terms of the statistical property generation as they did in the simulations. GRAMMAR found a few QTNs with high false-negative errors, whereas BOLT-LMM produced more QTNs than most other competing methods, with the exception of Optim-GRAMMAR with joint analysis, but had higher false-positive errors. With ideal GC, GRAMMAR-Gamma identified the same number of QTNs as Optim-GRAMMAR. We have described our findings of Optim-GRAMMAR and FaST-LMM simulations in the subsequent text.

The Q-Q profiles showed the GCs optimized to be very close to 1.0 by Optim-GRAMMAR. When joint association analyses are used in Optim-GRAMMAR, we found the QTNs in 26 of 109 phenotypes of HS mice, while and 35 of 36 phenotypes of CFW mice. However, FaST-LMM did not find any QTN in 28 of these 35 traits of CFW mice. Under the optimized GC, Optim-GRAMMAR identified 8 and 183, respectively, more the QTNs with joint analyses than FaST-LMM did in the HS mouse and CFW mouse. Even with a test at once in the HS mouse, Optim-GRAMMAR could cover 96% of the QTNs obtained with FaST-LMM. Especially in the CFW mouse, Optim-GRAMMAR completely covered the QTNs detected with FaST-LMM.

Finally, we applied the Optim-GRAMMAR to jointly analyze the QTNs for the flowering time in maize and test the performance by using 5 000 sampling SNPs to estimate GRM. By using entire markers, Optim-GRAMMAR detected 8 QTNs distributed on the chromosomes 1, 2, 3, 8, and 10, whereas could also find the same QTNs and an additional QTN detected on chromosome 7 by using the sampling SNPs to calculate GRM. With a test at once, Optim-GRAMMAR and FaST-LMM identified five the isolated QTNs. Of the detected QTNs, there were three identical pairs and one different pair located on chromosome 2 for Optim-GRAMMAR and chromosome 6 for FaST-LMM. In addition, the two Optim-GRAMMARs that use entire and sampling markers to estimate GRM took 28.21s and 25.37s, respectively, to implement 5-6 iterations of association tests in PLINK 2.0^29^. While, GRAMMAR and GRAMMAR-Gamma consumed 2.073 and 3.637 min, respectively, to do one iteration of the association tests. Additionally, FaST-LMM ran the Single-Runking ^30^ 32.147 min and BOLT-LMM ran the BOLT ^8^ 166.448 min for the association analysis.

## Discussion

In solving the GBLUP equations, the GBVs are underestimated by selecting the polygenic heritability instead of the genomic heritability of traits, which better approximates the minor polygenic effects except for major markers. In next association tests, only major effects are remained in residual phenotypes that polygenic effects are removed, so that GRAMMAR produces the minimum false-negative rate by administering a GC of 1.0, ensuring the same statistical power to detect QTNs as the exact mixed model association analysis. Joint analysis for multiple QTN candidates chosen through multiple testing further improves the statistical power as it takes into consideration the linkage disequilibrium among candidate markers.

Genome-wide mixed model association analysis is generally implemented in three steps ^31^: 1) building the GRM, 2) estimating variance components or polygenic heritability, and 3) computing association statistics for each SNP. The Optim-GRAMMAR algorithm proposed in our study inherits the lowest computing complexity of GRAMMAR for the association tests. Similar to the extremely fast BOLT-LMM that estimates the variance components with Monte Carlo REML ^11,12^, the Optim-GRAMMAR solves GBLUP equations multiple times to estimate the approximate polygenic effects through genomic heritability. However, our computing times are not greater than that of BOLT-LMM, especially for complex population structures. With the markers selected randomly across the whole genome for the calculation of the GRM, our proposed method retains a statistical power similar to that of the exact mixed model association analysis. For genomic dataset containing *m* SNPs genotyped on *n* individuals, Optim-GRAMMAR consumes only a *O*(*mn*^2^) computing time to build relationship matrices and *O*(*mn*) for association tests. When analyzing large-scale populations, we can solve the effects of *m*_0_, the sampled markers 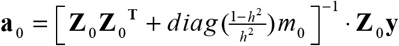, and given heritability *h*^2^, using ridge regression ^32^ and then estimate the GBVs as **Z_0_a_0_.** These reduce the computing time required to build information matrices to 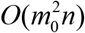, as seen in FaST-LMM-Select ^10^, so that the computing complexity of Optim-GRAMMAR becomes linear on population size. For the simulated 8 million SNPs on 400,000 individuals, Optim-GRAMMAR required 16.95 min, 2.51min of which to optimize GCs by using 5,000 and 20,000 SNPs to evaluate GRM and GCs, respectively, which ran as fast as fastGWA^27^ almost in the similar memory footprints. A user friendly Optim-GRAMMAR software was developed, which is freely available at https://github.com/RunKingProgram/Optim-GRAMMAR.

For improving statistical power, in general, joint association analysis for multiple QTN candidates with stepwise regression generated a higher estimating robustness and computing efficiency than the Gaussian mixture-model association analysis ^33-35^ of BOLT-LMM. Especially, the optimized residuals and heritability are not required to update in joint association analysis. Without a need to directly estimate the genomic heritability with a nonlinear solution, such as the penalized quasi-likelihood ^36^, our Optim-GRAMMAR can be easily extended to handle binary disease traits, greatly simplifying genome-wide generalized mixed model association analysis.

## Acknowledgements

The research is financially supported by the National Natural Science Foundations of China (32072726) and the Special Scientific Research Funds for Central Non-profit Institutes, Chinese Academy of Fishery Sciences (2019A001).

## Online Methods

### Genomic regression model

For mapping QTNs, quantitative traits were recorded and *m* high throughput SNPs genotyped for *n* individuals. These markers were distinguished from major and common alleles based on the magnitude of effects of the markers on quantitative traits. We described the relationship between all markers and the phenotypes using the following genomic regression model:

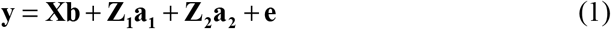

Where, **Xb** is representative of the fixed effect terms, such as population structure (stratification), sex, and age. **a_1_** represents the large genetic effects of the *q* markers on phenotype, and **a_2_** represents the minor or zero effects of the *m*-*q* markers on phenotype. The remaining variables represent the following: **X**, design matrices of fixed effects; **b**, **Z_1_**, and **Z_2_**, are indicator variables of SNP genotypes, which are generally coded as −1, 0, and 1 (or 0, 1 and 2) for the three genotypes AA, AB, and BB, respectively; **e**, residual error. The residual error is characterized as 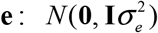, with residual variance 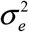 and **I** identity matrix.

### Statistical inference

Prior to estimating parameter, the indicator variables of SNP genotypes are generally centred and standardized to unit variance ^1,2^. Specifying **a**_2_ as a multivariate normal distribution 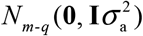, where 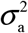 is a minor genetic variance, we obtain phenotypic variance-covariance matrix:

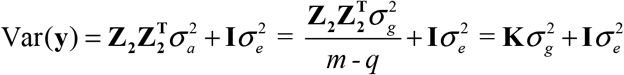

where **K** is the GRM between pairwise individuals calculated by using the minor or no effect SNPs and 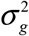 is polygenic variance 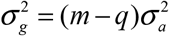. Thus, we transform genomic regression model to the LMM:

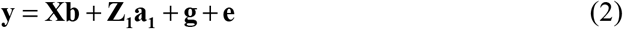

Where, **g** = **Z_2_a_2_** represents random polygenic effects and satisfies the assumption that 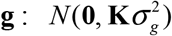

Within the framework of the GRAMMAR algorithm, we solve genomic mixed model (2) using two-stage regression method. The first stage is to estimate polygenic effects with the null model:

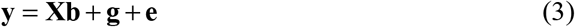

Thus, a GBLUP equations are constructed as:

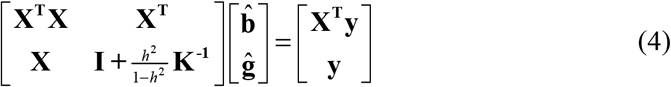

where *h*^2^ is the estimated genomic heritability. At the same time, we estimate phenotypic residuals as:

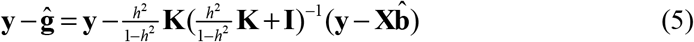

The second stage is to statistically infer association of each SNP with phenotypic residuals with simple regression model:

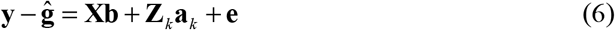

for the *k^th^* tested SNP.

It should be noted that at the first stage, we estimate the GBVs rather than polygenic effects. in addition of polygenic effects, there are a part of QTN effects in the GBVs. Moving partial QTN effects from phenotypic residuals makes genome-wide test statistics to be deflated at the second stage, which yields false-negative error in detecting QTNs.

### Optimization for GC

In the equations (4), estimation of polygenic effects mainly depends on genomic heritability. Accordingly, we can regulate downward genomic heritability to approximately estimate polygenic effects, until optimizing GC infinitely close to 1.0. In summary, we implement the optimized GRAMMAR by GC in following steps:

1. Initialize *h*^2^ to 0;
2. Estimate phenotypic residuals 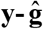 with equation (4) and (5);
3. Statistically infer the genetic effect for each SNP with chi-squared testing:

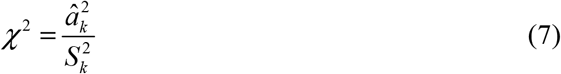

where 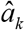 and 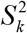 are the estimated SNP effect and its standard deviation with the model (6). The statistic *χ*^2^: *χ*^2^ (1) with 1 degree of freedom.
4. Calculate the genome-wide chi-squared mean and/or statistical probability for each SNP;
5. Plot the Quantile-Quantile (Q-Q) profile for the genome-wide statistical probabilities.
6. Update the *h*^2^ between 0 and the genomic heritability using Brent’s method ^3^;
7. Repeat steps (2)-(6) until the genome-wide chi-squared mean reaches 1^+^ or yields a satisfactory Q-Q plot.

In each iteration, only GBLUP equations are solved, this greatly simplifies the mixed model association analysis.

### Joint association analysis

For the residuals 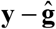 obtained with the Optim-GRAMMAR algorithm, we jointly analyzed multiple QTN candidates to improve the statistical power. At less than the Bonferroni-corrected criteria ^4^, a lot of the SNPs with relatively low linkage disequilibria were chosen as QTN candidates. In principle, the number of QTN candidates would not be greater than the population size. We adopted a backward regression to optimize the multiple linear regression model as follows:

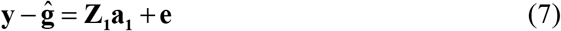

Given the Bonferroni-corrected significance level, the significant QTN effects were filtered in a stepwise manner according to the test statistic (7), allowing for the identification of the QTNs.

## Simulations

Two representational genomic datasets in human and maize were used to simulate the adaptability of Optim-GRAMMAR to population structure. Maize population ^5^ has a more complex structure than the human population ^6^. We extracted 300,000 SNPs for both people (n=3000) and maize (n=2640) through higher quality control. We distributed randomly simulated QTNs over these SNPs. Additive effects of the QTNs were characterized from a gamma distribution with a shape (1.66) and scale (0.4) such that there were few with large effects and more with minor effects in the simulated QTNs. Phenotypes were obtained by summing the genotypic effects of all the simulated QTNs and residual errors. In the sampling of the residual errors from a normal distribution with zero expectation, residual variance was regulated by the given genomic heritability of traits.

In addition to the population structure, the number of QTNs, genomic heritability and sampling numbers of SNPs were also considered as experimental factors in the simulations. We simulated phenotypes controlled by 40, 200, and 1000 QTNs with varied levels of heritability (low, 0.2; moderate, 0.5; and high, 0.8). The GRMs and GCs were calculated based on clusters of 3000, 5000, 10 000, 15 000, 20 000, and 25 000 SNPs drawn from entire genomic markers. Simulation experiments were performed to investigate the following: (1) statistical properties of Optim-GRAMMAR under different combinations of genomic heritability and number of QTNs simulated; and (2) calculations of GRMs with the sample markers for the above-mentioned simulated phenotypes controlled by 200 QTNs. In both simulation experiments, we compared Optim-GRAMMAR, a test at once, with FaST-LMM, GRAMMAR, GRAMMAR-Gamma, and BOLT-LMM, using Q-Q profiles or GCs and ROC profiles. In the first simulation experiment, we performed joint association analysis of Optim-GRAMMAR to improve the statistical power.

Under good GC (very close to 1.0, in general), the ROC profiles can be plotted by statistical powers to detect QTNs relative to given a series of Type I errors. Statistical powers are defined as the percentage of identified QTNs that have the maximum test statistic among their 20 closest neighbors over the total number of simulated QTNs. Simulations are repeated 50 times, with the average results recorded. Note that the positions and effects of QTNs simulated vary in each repeated experiment.

## Real data

We applied the Optim-GRAMMAR algorithm to analyze datasets collected from three different populations: 1) heterogeneous stock mice (HS mice, n=1,940) with 12,226 SNPs genotyped and 123 phenotypes recorded ^7^, 2) Carworth Farms White outbred mice (CFW mice, n=1149) with the 92,734 SNPs genotyped and the 973 physiological traits observed ^8^; and 3) maize inbred lines (n=2,279) with 681,258 SNPs genotyped and flowering time (measured as days to silk) mapped ^9^ (URL:http://www.panzea.org/!#genotypes/cctl). All real data analyses were performed on a computing platform with 2.60 GHz Intel(R) Xeon(R) 40 CPUs E5-2660 v3, and 512 GB memory.

## References

1. Schaeffer, L.R. The Animal Models, (University of Guelph, Guelph, 2019).

2. Yu, J.M. et al. A unified mixed-model method for association mapping that accounts for multiple levels of relatedness. Nature Genetics 38, 203–208 (2006).

3. Price, A.L., Zaitlen, N.A., David, R. & Nick, P. New approaches to population stratification in genome-wide association studies. Nature Reviews Genetics 11, 459–463 (2010).

4. Kariya, T. & Kurata, H. Generalized Least Squares, (John Wiley & Sons, Chichester, UK, 2004).

5. Kang, H.M. et al. Efficient control of population structure in model organism association mapping. Genetics 178, 1709–1723 (2008).

6. Lippert, C. et al. FaST linear mixed models for genome-wide association studies. Nature Methods 8, 833–835 (2011).

7. Zhou, X. & Stephens, M. Genome-wide efficient mixed-model analysis for association studies. Nature Genetics 44, 821–824 (2012).

8. Loh, P.R. et al. Efficient Bayesian mixed-model analysis increases association power in large cohorts. Nature Genetics 47, 284–290 (2015).

9. Zhang, Z.W. et al. Mixed linear model approach adapted for genome-wide association studies. Nature Genetics 42, 355–360 (2010).

10. Jennifer, L. et al. Improved linear mixed models for genome-wide association studies. Nature Methods 9, 525–526 (2012).

11. García-Cortés, L.A. et al. Variance component estimation by resampling. Journal of Animal Breeding and Genetics 109, 358–363 (1992).

12. Matilainen, K., Mantysaari, E.A., Lidauer, M.H., Stranden, I. & Thompson, R. Employing a Monte Carlo algorithm in Newton-type methods for restricted maximum likelihood estimation of genetic parameters. PLoS One 8, e80821 (2013).

13. Haseman, J.K. & Elston, R.C. The investigation of linkage between a quantitative trait and a marker locus. Behavior Genetics 2, 3–19 (1972).

14. Chen, G.-B. Estimating heritability of complex traits from genome-wide association studies using IBS-based Haseman-Elston regression. Frontiers in Genetics 5, 1–14 (2014).

15. Patterson, H.D. & Thompson, R. Recovery of inter-block information when block sizes are unequal. Biometrika 58, 545–554 (1971).

16. Reverter, A., Golden, B.L., Bourdon, R.M. & Brinks, J.S. Method R variance components procedure: application on the simple breeding value model. J Anim Sci 72, 2247–53 (1994).

17. Coleman, J. Genomic Testing and Method R Variance Components Theory of Dairy Cattle. (2012).

18. Vanraden, P.M. et al. Invited review: reliability of genomic predictions for North American Holstein bulls. Journal of Dairy Science 92, 16–24 (2009).

19. Aulchenko, Y.S., de Koning, D.J. & Haley, C. Genomewide rapid association using mixed model and regression: a fast and simple method for genomewide pedigree-based quantitative trait loci association analysis. Genetics 177, 577–585 (2007).

20. Kang, H.M. et al. Variance component model to account for sample structure in genome-wide association studies. Nature Genetics 42, 348–354 (2010).

21. Svishcheva, G.R., Axenovich, T.I., Belonogova, N.M., van Duijn, C.M. & Aulchenko, Y.S. Rapid variance components-based method for whole-genome association analysis. Nature Genetics 44, 1166–1170 (2012).

22. Meuwissen, T.H., Hayes, B.J. & Goddard, M.E. Prediction of total genetic value using genome-wide dense marker maps. Genetics 157, 1819–1829 (2001).

23. Gianola, D. Priors in whole-genome regression: the bayesian alphabet returns. Genetics 194, 573–596 (2013).

24. Friedman, J., Hastie, T. & Tibshirani, R. Regularization Paths for Generalized Linear Models via Coordinate Descent. Journal of Statistical Software 33, 1–22 (2010).

25. Henderson, C.R. Applications of linear models in animal breeding, (University of Guelph Guelph, 1984).

26. Vanraden, P.M. Efficient methods to compute genomic predictions. Journal of Dairy Science 91, 4414–4423 (2008).

27. Jiang, L. et al. A resource-efficient tool for mixed model association analysis of large-scale data. Nature Genetics 51, 1749–1755 (2019).

28. Yang, J. et al. Common SNPs explain a large proportion of the heritability for human height. Nature Genetics 42, 565–569 (2010).

29. Purcell, S. et al. PLINK: a tool set for whole-genome association and population-based linkage analyses. American Journal of Human Genetics 81, 559–575 (2007).

30. Gao, J., Zhou, X., Hao, Z., Jiang, L. & Yang, R. Genome-wide barebones regression scan for mixed-model association analysis. Theor Appl Genet (2019).

31. Yang, J., Zaitlen, N.A., Goddard, M.E., Visscher, P.M. & Price, A.L. Advantages and pitfalls in the application of mixed-model association methods. Nature Genetics 46, 100–106 (2014).

32. Hoerl, A.E. & Kennard, R.W. Ridge Regression: Biased Estimation for Nonorthogonal Problems. Technometrics 12, 55–67 (1970).

33. Logsdon, B.A., Hoffman, G.E. & Mezey, J.G. A variational Bayes algorithm for fast and accurate multiple locus genome-wide association analysis. BMC Bioinformatics 11, 58 (2010).

34. Logsdon, B.A., Carty, C.L., Reiner, A.P., Dai, J.Y. & Kooperberg, C. A novel variational Bayes multiple locus Z-statistic for genome-wide association studies with Bayesian model averaging. Bioinformatics 28, 1738–44 (2012).

35. Stephens, M. & Balding, D.J. Bayesian statistical methods for genetic association studies. Nat Rev Genet 10, 681–90 (2009).

36. Breslow, N.E. & Clayton, D.G. Approximate inference in generalized linear mixed models. Journal of the American statistical Association 88, 9–25 (1993).

## Method References

1. Hayes, B.J., Visscher, P.M. & Goddard, M.E. Increased accuracy of artificial selection by using the realized relationship matrix. Genet Res (Camb) 91, 47–60 (2009).

2. Meuwissen, T.H., Solberg, T.R., Shepherd, R. & Woolliams, J.A. A fast algorithm for BayesB type of prediction of genome-wide estimates of genetic value. Genetics Selection Evolution 41, 2 (2009).

3. Brent, R.P. Algorithms for minimization without derivatives, (Prentice-Hall, New Jersey, 1973).

4. Hochberg, Y. & Tamhane, A.C. Multiple Comparison Procedures, (John Wiley & Sons, Inc., New York, 1987).

5. Consortium, W.T.C.C. Genome-wide association study of 14,000 cases of seven common diseases and 3,000 shared controls. Nature 447, 661–78 (2007).

6. Romay, M.C. et al. Comprehensive genotyping of the USA national maize inbred seed bank. Genome biology 14, R55 (2013).

7. Solberg, L.C. et al. Genome-wide genetic association of complex traits in heterogeneous stock mice. Nature Genetics 38, 879–887 (2006).

8. Parker, C.C. et al. Genome-wide association study of behavioral, physiological and gene expression traits in outbred CFW mice. Nature genetics 48, 919–926 (2016).

9. Romay, M.C. et al. Comprehensive genotyping of the USA national maize inbred seed bank. Genome Biol 14, R55 (2013).

